# The R package divDyn for quantifying diversity dynamics using fossil sampling data

**DOI:** 10.1101/423780

**Authors:** Kocsis Á. T., Reddin C. J., Alroy J., Kiessling W.

## Abstract

1. Unbiased time series of diversity dynamics are vital for quantifying the grand history of life. Applications include identifying ancient mass extinctions and inferring both biotic and abiotic controls on diversification rates.
2. We introduce divDyn, a new R package that facilitates the calculation of taxonomic richness, extinction and origination rates from time-binned fossil sampling data. State-of-the-art sampling completeness metrics, counting protocols, and sampling standardisation functions permit the reconstruction of biologically meaningful time series. Additional functions permit the partitioning of turnover rates by trait and environmental affinity.
3. We display Phanerozoic-scale diversity dynamics of marine invertebrates using the divDyn package. Using the core function and standard subsampling options, we revisit earlier assessments of declining extinction and origination rates over time and of equilibrial diversity dynamics to assess their methodological dependency.
4. The modular and fast implementation of published methods ensures traceability, reproducibility, and comparability of future studies.

## Introduction

Reconstructions of global diversity (Sepkoski *et al.* 1981; Alroy *et al.* 2001; Alroy *et al.* 2008; Alroy 2010a) as well as extinction and origination rates (Newell 1952; Carr & Kitchell 1980; Raup & Sepkoski 1982; Sepkoski 1993; Benton 1995; Foote 2000; Bambach, Knoll & Wang 2004; Alroy 2008; Alroy 2014; Alroy 2015) have led to the recognition of ancient mass extinctions (Raup & Sepkoski 1982; Alroy 2008) and major insights about the interplay of evolutionary crisis and recovery. All of these results depend on the reconstruction of time series that characterise changes in the global biosphere. Calculating these time series from an incomplete fossil record is a fundamental task, as they serve as the basis for the statistical testing of grand questions in macroevolution.

Early studies largely relied on compendia of stratigraphic ranges, that is, they derived diversity metrics from overlapping durations of taxa (e.g. Sepkoski 1984). Since the advent of the Paleobiology Database (PaleoDB, https://paleobiodb.org) diversity dynamics have largely been inferred from large occurrence datasets that incorporate hundreds of thousands of items. Occurrence data allow for alternative counting methods and sampling standardisation, but implementing these methods in scripting languages is time-consuming and can be challenging for students and researchers with little programming experience. The algorithmic implementation of some procedures and the multiple steps of data filtering also permit considerable analytical freedom, which potentially compromises the comparability and traceability of results. Using a standardised toolkit will facilitate a fast and consistent workflow and allows researchers to focus on scientific questions rather than losing time with repeated implementation of established methods in computer scripts.

Here we present the R (2018) package divDyn, which facilitates the calculation of diversity dynamics from fossil occurrence datasets along with additional methods of palaeobiological inference. Our purpose is to establish a transparent, traceable and modular workflow from data acquisition to the calculation of biologically meaningful diversity metrics. The primary application of the package is expected to be for data from the Paleobiology Database (PaleoDB, https://palebiodb.org/), the largest fossil occurrence dataset, which serves as a standard for palaeontological data analyses. However, any dataset for which diversity metrics are to be assessed in temporal or spatial intervals can also be processed in divDyn. The PaleoDB, Neptune (Lazarus 1994) and AMMON (Korn & Ilg 2007) databases are good examples of alternative datasets. To demonstrate the advantages of divDyn, we revisit results from an earlier study on Phanerozoic-scale diversity dynamics (Alroy 2008).

## Features

### Calculation of Time Series

To prepare an occurrence dataset for analysis in divDyn, it must be formatted as a table with each row representing a single occurrence of a taxon. Occurrences should be assigned to discrete time intervals (bins). Alternatively, values in a continuous time-related dimension (e.g. years before present, or meters in a section) can be used that will be translated to discreet bins by the package.

The core function `divDyn()` calculates taxonomic richness, extinction and origination rate estimates using the presence-absence patterns of the bin-taxon matrix implied by the input dataset. It calculates a large suite of diversity metrics and rates in one go, ranging from classical sampled-in-bin richness to the recent second-for-third proportions (Alroy 2015, Table 1).

**Table 1.**
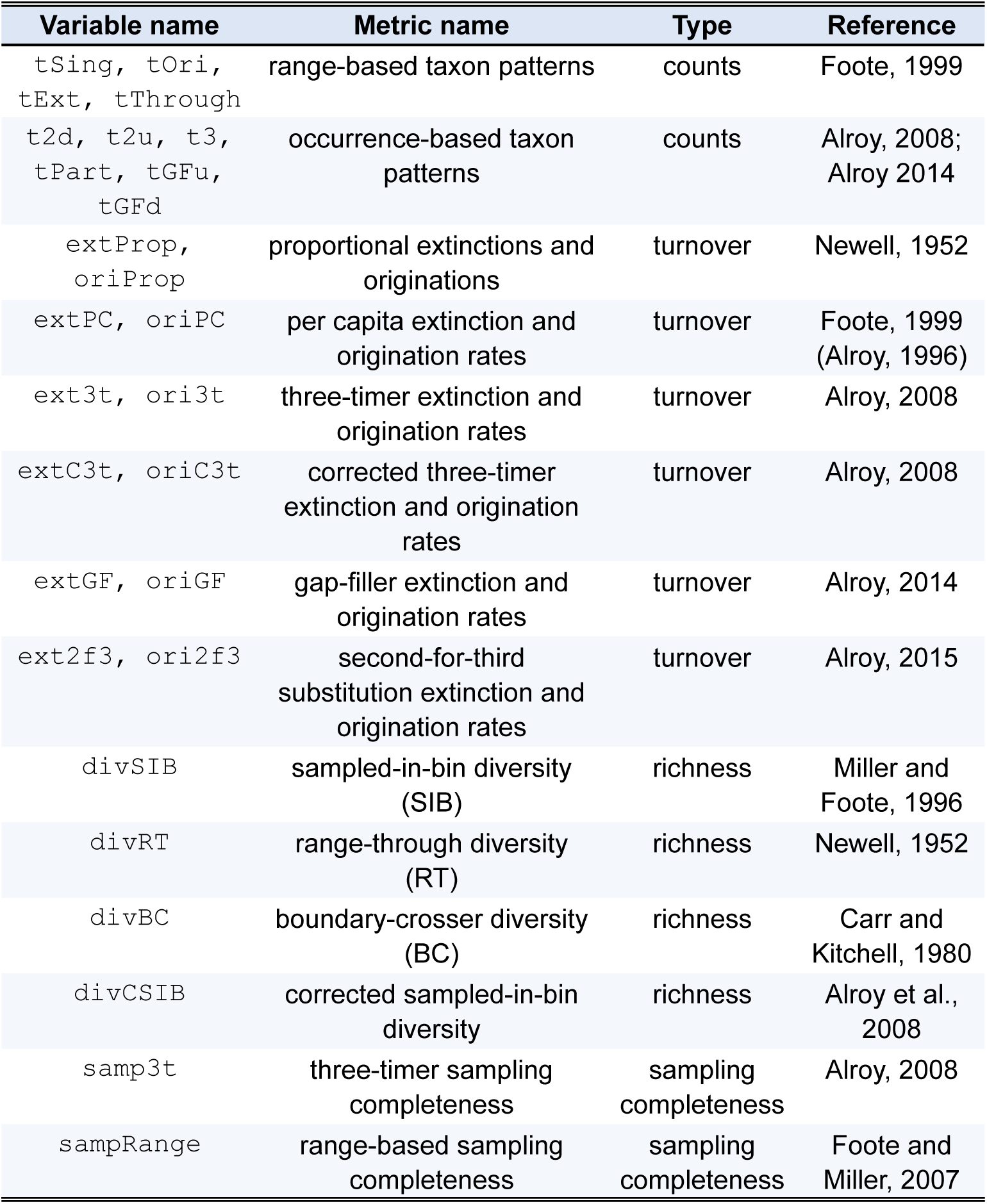
List of some of the variables output by the main function of the package.

### Sampling Standardisation Procedures

Sampling standardisation is a useful tool for smoothing out the effects of changing sampling intensity on the available data. Subsampling, the most common group of methods in this field of research, uses interpolation to answer the general question ‘What would the results look like if fewer data were available?’. The oldest and most widely used subsampling method is rarefaction (Sanders 1968; Miller & Foote 1996) but there are many others (Alroy *et al.* 2001; Bush, Markey & Marshall 2004; Alroy 2010a; Chao & Jost 2012). More implementations of these methods are available for estimating richness from an incomplete sample (Hammer, Harper & Ryan 2001; Hsieh, Ma & Chao 2016), but their application is more complicated in the context of time series reconstruction (Fig. 1), where information combined from multiple samples is to be extracted.

**Figure 1.**
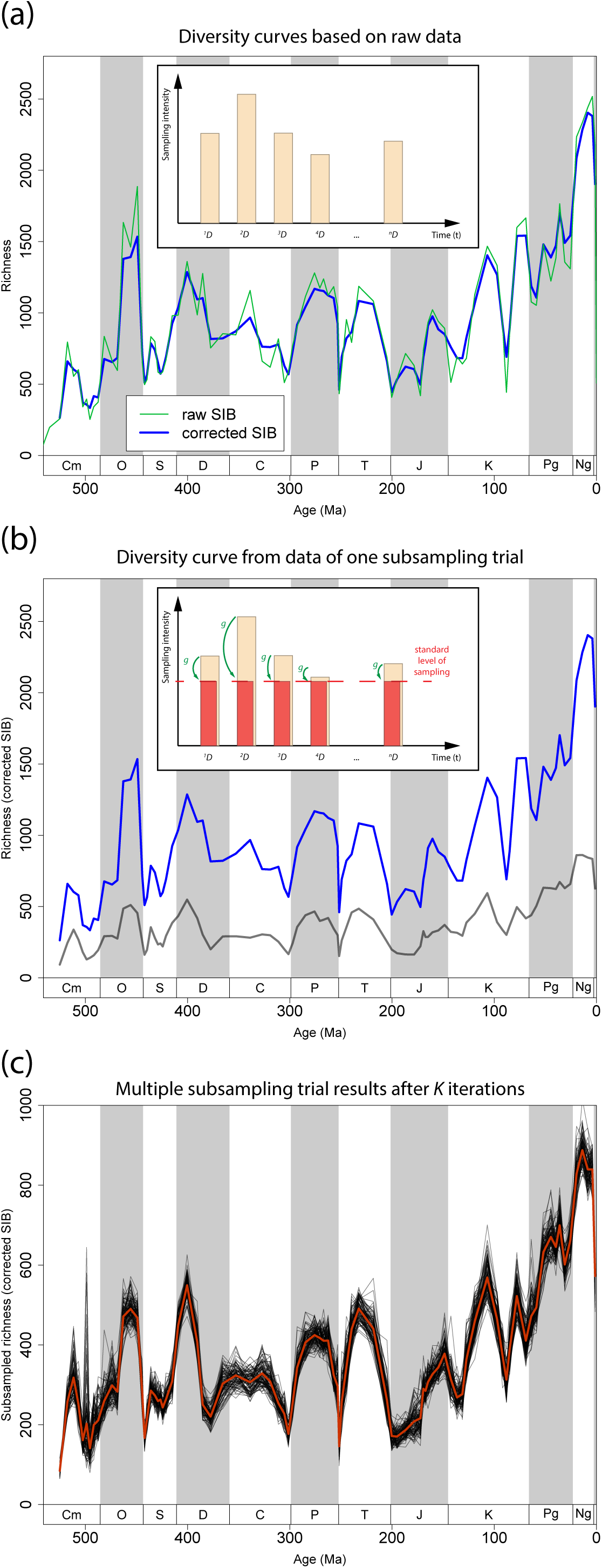
Demonstration of the procedures implemented in the subsampling wrapper function. (a) Calculation of raw results, (b) calculation of results from a single subsampling trial and (c) multiple trial results and averaging. The curves show genus richness from the Phanerozoic example dataset, standardized with SQS at the stratigraphic resolution of geologic stages. The number of iterations (*K*) was 100, the quorum for SQS is 0.7.

We formalized the subsampling process in the wrapper function `subsample()`. The arguments of the process are *D*, g, *f*, *K*, α_f_ and α_g_, where *D* is the source dataset, *g* is the subsampling function, *f* is the user-supplied function, *K* is the number of iterations, and α_f_ and α_g_ are sets of additional function arguments for *f* and *g*, respectively. The desired result, based on all sampled data, is just a function of the data and its own arguments

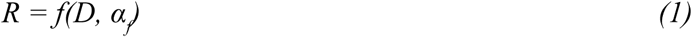

Given that the dataset is divisible into finite subsets, such as time bin-specific parts

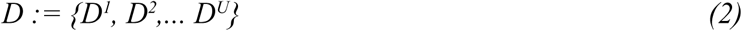

each bin-specific part has an abstract subset that features the same desired sampling characteristics (for instance, intensity), thus defining a set of data

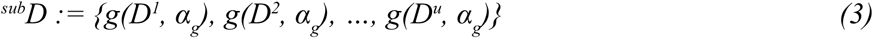

With this *^sub^D* abstract representation of the data, the desired results (*^sub^R*) would be just

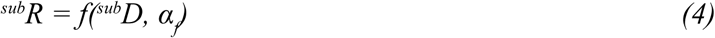

This can be approximated with Monte Carlo simulations, by generating multiple, randomly subsampled instances of *^sub^D_k_* (subsampled dataset in trial *k* out of *K*) and allowing the naturally emerging variation to propagate to *^sub^R_k_* (subsampled result in trial *k*). Thus, the simulation will lead to a vector of possible results.

From the point of implementation, *f* can be any function that is applicable to the original occurrence dataset *D* and the class of *^sub^R_k_* can be of any sort, such as vectors (time series) and complex structures, if they can be vectorized in the form of a list-class R object. In the case of primitive trial results (vectors and matrices), the averaging of time series can be automated. The current package version features Classical Rarefaction (Sanders 1968; Miller & Foote 1996), Occurrence-Weighted By-List Subsampling (O^x^W, Alroy *et al.* 2001), and Shareholder Quorum Subsampling (SQS, Alroy 2010a).

### Additional Functionality

To facilitate the stratigraphic assignment of collections, we compiled tables using the dynamic timescale interpreter of Fossilworks (http://fossilworks.org/) that links entries to major geochronological intervals of two predefined timescales (the level of geological stages and the 10-million-year time scale). The package also includes additional tables to categorise the downloaded occurrences in terms of bathymetry (shallow, deep), substrate (siliciclastic vs. carbonate) and reef vs. non-reef environments.

Occurrence patterns of taxa allow the inference of preferred environments, which we can use to compare diversity dynamics between groups with different affinities such as tropical vs. extra-tropical or reef vs. non-reef (Alroy 2010a; Kiessling, Simpson & Foote 2010). As fossil sampling is biased and heterogeneous, we implemented methods to calculate likely environmental preferences with sampling patterns taken explicitly into account (Kiessling & Aberhan 2007), as well as test for selectivity of extinctions (Kiessling & Kocsis 2015).

The `divDyn()` function creates output in discretised time intervals. Results can be visualized effectively with the additional plotting functions that display the used time scale (`tsplot()`), stratigraphic ranges (`ranges()`), changes in composition (`parts()`) or distributions of values in time series (`shades()`).

The basic functionality of the package is elaborated in the accompanying vignette (Handout to the R package divDyn v0.6.0 for diversity dynamics from fossil occurrence data), with an example dataset of Scleractinian corals used by Kiessling and Kocsis (2015).

## Example Application: Phanerozoic-Scale Diversity Dynamics of Marine Animals

Tracing diversity through the entire Phanerozoic (the last 541 million years of Earth history) has been the focus of palaeobiological research ever since the first global diversity curves were published (Newell 1952). Some patterns have been surprisingly robust to scientific scrutiny, whereas others are more volatile. For example, the temporal decline of turnover rates has largely gone unchallenged since its first observation (Raup & Sepkoski 1982), whereas the original ‘Big Five’ mass extinction events of Raup and Sepkoski (1982) have been repeatedly revisited, with different conclusions (Bambach, Knoll & Wang 2004; Alroy 2008). Much discussion has focused on the dramatic rise of marine biodiversity over the last 100 myr, which is evident in older compilations (Valentine 1970; Sepkoski 1993) but much less so in sampling-standardised analyses (Alroy *et al.* 2008). Not yet formally contested are Alroy’s (2008, 2010b) analyses of the temporal relationship between diversity and rates. If these results are robust at different temporal resolutions, they strongly argue for equilibrial, diversity-dependent diversity dynamics (Sepkoski 1978; Sepkoski 1984; Alroy 1996) through biotic interactions. With the continuous expansion of both fossil occurrence data sets and the toolkit to analyse them, it is necessary to re-evaluate such scientific outcomes on a periodical basis. The objective of this case study is to assess the robustness of previous results in the face of the increase in the number of fossil occurrences and the number of choices we face when we express diversity dynamics over deep time.

### Data Processing and Applied Methods

The analyses presented in this section can be reproduced with the second vignette accompanying the package (Phanerozoic-scale global marine biodiversity analysis with the R package divDyn v0.6; http://github.com/adamkocsis/ddPhanero). The occurrences used here were downloaded on 14 September 2018, including all entries between the Ediacaran-Holocene interval. The data were filtered to marine taxa and binned to geological stages as well the often used 10 myr bins (Alroy et al. 2008). As the procedural treatment of stages in the Cambrian and Ordovician systems was influenced by considerable stratigraphic error, they were resolved using biozone and formation entries (Ordovician), and with data from previous analyses (Cambrian, Na & Kiessling 2015). In keeping with related literature (e.g. Sepkoski 1997), all analyses were carried out at the genus level. However, species-level analyses can be conducted with exactly the same procedures.

We computed diversity dynamics at both stratigraphic resolutions (stages and 10 myr), with three different treatments of the data (raw, CR and SQS). Four different rate metrics were applied: per capita rates (Foote 1999, used most frequently in studies), corrected three-timer rates (Alroy 2008), gap-filler equations (Alroy 2014), and second-for-third substitution rates of Alroy (2015). This resulted in 24 different sets (2 timescales × 3 data treatments × 4 rate metrics) of richness, origination and extinction rate series, each affected in a different way by the distorting effects of incomplete, heterogeneous sampling and estimation error. Simple correlation and normality tests were applied to each of those sets to check whether previous results are robust to varying protocols.

As indicated by Foote (2005), most taxonomic turnover was probably pulsed, likely at stage boundaries. This assertion is supported by the infrequent correlations between interval durations and rate values when the time dimension is excluded from the rate equations (Table 2 and Alroy 2008). For the analysis of distributions, outliers and cross correlations, we detrended the rates and the richness by applying LOESS with AICc-based smoothing parameters (Wang 2010) to describe long-term variation (Bambach, Knoll & Wang 2004). Mass extinctions are defined as statistical outliers (using boxplot statistics) after the LOESS trend has been removed.

**Table 2.** Results of method-specific outcomes of the time series calculations for 10 myr bins (a) and stages (b). Unless otherwise indicated, values are estimates of correlation coefficients. Significant results of Shapiro-Wilk tests indicates deviation from log-normality. Formatting indicates significance (except for binary information): insignificant values are not reported, regular entries indicate 0.05 ≥ *p* > 0.01, **bold** entries 0.01 ≥ *p* > 0.001 and **<u>underscored bold</u>** entries denote *p* ≤ 0.001. Rate metrics are abbreviated: PC = the per capita rates (Foote 1999), C3t = corrected three-timer rates (Alroy 2008), GF = gap-filler rates (Alroy, 2014) and 2f3 = second-for-third-substitution rates (Alroy, 2015).

| (a) |  | 10 MYR TIME SCALE |  |  |  |  |  |  |  |  |  |  |  |
| --- | --- | --- | --- | --- | --- | --- | --- | --- | --- | --- | --- | --- | --- |
|  |  | raw |  |  |  | CR |  |  |  | SQS |  |  |  |
|  |  | PC | C3t | GF | 2f3 | PC | C3t | GF | 2f3 | PC | C3t | GF | 2f3 |
| Pulsed/<br>continuous | ext. rates with durations |  |  |  |  |  |  |  |  |  |  |  |  |
|  | norm. ext. rates with durations | <u>-0.49</u> | <u>-0.44</u> | -0.34 | -0.3 | <u>-0.56</u> | <u>-0.58</u> | <u>-0.55</u> | <u>-0.55</u> | <u>-0.47</u> | <u>-0.46</u> | -0.32 | -0.34 |
|  | orig. rates with durations |  |  | 0.32 |  |  |  |  |  |  |  |  |  |
|  | norm. orig. rates with durations | -0.37 | <u>-0.42</u> |  |  | -0.36 | <u>-0.4</u> |  |  |  | -0.35 |  |  |
| Declines | extinctions | <u>0.55</u> | <u>0.58</u> | <u>0.61</u> | <u>0.51</u> | <u>0.55</u> | <u>0.61</u> | <u>0.62</u> | <u>0.57</u> | <u>0.55</u> | <u>0.61</u> | <u>0.65</u> | <u>0.57</u> |
|  | post-Ordovician extinctions |  | 0.38 | <u>0.42</u> |  |  | <u>0.45</u> | <u>0.44</u> | 0.35 |  | 0.41 | <u>0.44</u> |  |
|  | originations | <u>0.5</u> | <u>0.45</u> | <u>0.5</u> |  | <u>0.55</u> | <u>0.43</u> | <u>0.52</u> | <u>0.41</u> | <u>0.53</u> | <u>0.43</u> | <u>0.53</u> |  |
|  | post-Ordovician originations |  |  |  |  |  |  |  |  |  |  |  |  |
| Mass extinctions | end-Permian ME | yes | yes | yes | yes | yes | yes | yes | yes | yes | yes | yes | yes |
|  | end-Triassic ME | yes | yes | yes | yes | yes | yes | yes | yes | yes | yes | yes | yes |
|  | end-Cretaceous ME | yes | yes | yes | yes | no | yes | yes | yes | no | yes | no | no |
|  | end-Permian is highest | yes | no | yes | yes | yes | yes | yes | yes | yes | yes | yes | yes |
|  | number of mass extinctions | 5 | 3 | 4 | 3 | 2 | 4 | 3 | 3 | 2 | 3 | 2 | 3 |
| Rate dist. | extinctions log-normal (p-values) |  |  |  |  |  | 0.04 |  |  |  |  |  |  |
|  | originations log-normal (p-values) |  |  |  | <u><0.001</u> |  |  |  |  |  |  | <u>0.01</u> | <u>0.01</u> |
| Equilibrial dyn. | origination and lagged diversity |  |  |  |  | 0.35 |  |  | 0.32 | <u>0.45</u> |  | 0.31 | 0.35 |
|  | diversity and lagged extinction | <u>0.45</u> |  |  |  | 0.35 |  |  |  | <u>0.48</u> |  |  |  |
|  | diversity and lagged origination |  | <u>0.55</u> |  |  |  | <u>0.67</u> | <u>0.48</u> | <u>0.42</u> |  | <u>0.54</u> |  |  |

| (b) |  | STAGE-LEVEL TIME SCALE |  |  |  |  |  |  |  |  |  |  |  |
| --- | --- | --- | --- | --- | --- | --- | --- | --- | --- | --- | --- | --- | --- |
|  |  | raw |  |  |  | CR |  |  |  | SQS |  |  |  |
|  |  | PC | C3t | GF | 2f3 | PC | C3t | GF | 2f3 | PC | C3t | GF | 2f3 |
| Pulsed/<br>continuous | ext. rates with durations | 0.23 |  | 0.22 | <u>0.28</u> |  |  |  | 0.22 |  |  |  | <u>0.27</u> |
|  | norm. ext. rates with durations | <u>-0.45</u> | <u>-0.35</u> | <u>-0.34</u> |  | <u>-0.64</u> | <u>-0.53</u> | <u>-0.58</u> | <u>-0.42</u> | <u>-0.54</u> | <u>-0.51</u> | <u>-0.46</u> | <u>-0.28</u> |
|  | orig. rates with durations | <u>0.35</u> |  | <u>0.32</u> | <u>0.33</u> |  |  |  |  | 0.24 |  | 0.24 | 0.23 |
|  | norm. orig. rates with durations | <u>-0.52</u> | <u>-0.51</u> | <u>-0.4</u> | <u>-0.31</u> | <u>-0.66</u> | <u>-0.58</u> | <u>-0.55</u> | <u>-0.51</u> | <u>-0.59</u> | <u>-0.5</u> | <u>-0.41</u> | <u>-0.39</u> |
| Declines | extinctions | <u>0.52</u> | <u>0.39</u> | <u>0.39</u> | <u>0.36</u> | <u>0.59</u> | <u>0.43</u> | <u>0.46</u> | <u>0.46</u> | <u>0.59</u> | <u>0.4</u> | <u>0.39</u> | <u>0.37</u> |
|  | post-Ordovician extinctions |  |  |  |  | 0.27 |  |  |  | 0.29 |  |  |  |
|  | originations | <u>0.44</u> | <u>0.36</u> | <u>0.39</u> | <u>0.35</u> | <u>0.58</u> | <u>0.39</u> | <u>0.43</u> | <u>0.43</u> | <u>0.52</u> | <u>0.36</u> | <u>0.39</u> | <u>0.38</u> |
|  | post-Ordovician originations |  |  |  |  | 0.24 |  |  |  |  |  |  |  |
| Mass extinctions | end-Permian ME | yes | yes | yes | yes | yes | yes | yes | yes | yes | yes | yes | yes |
|  | end-Triassic ME | yes | yes | yes | yes | yes | yes | yes | yes | yes | yes | yes | yes |
|  | end-Cretaceous ME | yes | yes | yes | yes | yes | yes | yes | yes | yes | yes | yes | yes |
|  | end-Permian is highest | yes | yes | yes | yes | yes | yes | yes | yes | yes | yes | yes | yes |
|  | number of mass extinctions | 4 | 3 | 5 | 4 | 3 | 3 | 3 | 4 | 3 | 3 | 6 | 3 |
| Rate dist. | extinctions log-normal (p-values) |  |  | 0.01 | <u><0.001</u> |  |  | <u><0.001</u> |  |  |  | <u><0.001</u> | <u><0.001</u> |
|  | originations log-normal (p-values) |  | 0.05 | <u><0.001</u> |  |  |  | <u><0.001</u> | <u><0.001</u> |  |  | <u><0.001</u> | 0.02 |
| Equilibrial dyn. | origination and lagged diversity | <u>0.39</u> | 0.3 |  |  | <u>0.34</u> |  |  |  | <u>0.52</u> | <u>0.34</u> | <u>0.34</u> | 0.28 |
|  | diversity and lagged extinction | <u>0.6</u> | <u>0.41</u> | <u>0.39</u> | <u>0.34</u> | <u>0.45</u> |  |  |  | <u>0.61</u> | 0.23 | <u>0.37</u> | <u>0.3</u> |
|  | diversity and lagged origination |  |  |  |  |  | <u>0.46</u> | <u>0.46</u> | <u>0.38</u> |  |  |  |  |

### Results

The first order patterns of the time series acquired with the different methods match very well, but the estimates for the individual time slices (Fig. 2) and thus the derived conclusions (Table 2) vary considerably. Although the decline of extinction and origination rates can be confirmed if the Cambrian and Ordovician is included in the dataset, the rates are unlikely to have featured a solid decline after the Ordovician period. All detrended extinction-rate series feature the latest Permian value as a mass extinction, which is consistently identified as the highest value in the series. The number of mass extinctions vary considerably. Among the ‘Big Five’ mass extinctions (Raup & Sepkoski 1982), the end-Triassic and the end-Cretaceous values also show up as outliers, as was indicated by Alroy (2008). The cross-correlations that indicate equilibrial dynamics appear to be unstable and are likely affected by the estimation error of the values in the time series.

**Figure 2.**
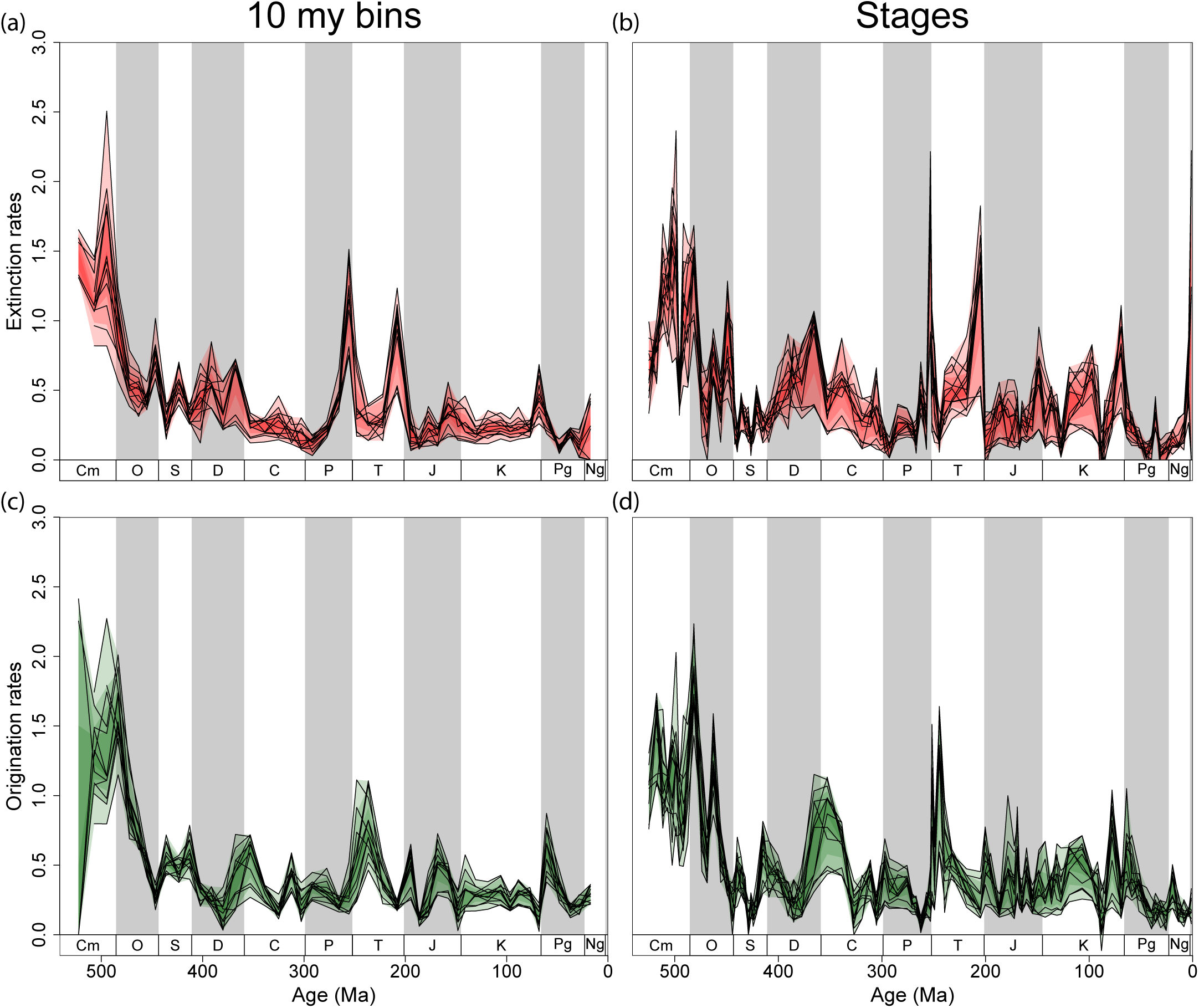
Genus-level Phanerozoic-scale extinction (a-b) and origination rates (c-d) calculated at the level of 10 million year bins and stratigraphic stages. Each panel features twelve (three treatment × four rate metrics) curves, either raw, CR (quotas are 4,800 for the 10my bins and 1,100 for the stages) or SQS-standardized (quorum is 0.7 for both) average per capita rates, corrected three-timer rates, gap-filler rates or second-for-third substitution rates.

### Discussion

Most major results derived from the Phanerozoic-scale analyses of the marine fossil animal record are robust after the additional 10 years of data entry and methodological development since Alroy (2008). Sepkoski (1993) found a similar pattern in analysing an older, range-based Phanerozoic diversity data set. One notable exception is the decline of rates through time, which receives varying support if the first two Phanerozoic periods are excluded. Likewise, changing data treatments either question equilibrial diversity dynamics or point to the possibility that carrying capacities change through time (Sepkoski 1984; Alroy 2010b; Marshall & Quental 2016).

Whichever the case, using a standard toolkit like divDyn enhances our ability to reproduce previous results and test the effect of added data or changing temporal resolution. We hope that our package will spur large-scale diversity analyses beyond the still small group of trained peers, such that results can be incorporated into broader evolutionary questions (Jablonski & Shubin 2015).

We intend to expand the set of output variables in the future, and the flexibility to write custom subsampling functions will be incorporated into divDyn. Our package utilises methods developed mostly by palaeobiologists. Simulations are currently being developed to compare results with other approaches such as PyRate (Silvestro, Salamin & Schnitzler 2014) or capture-mark-recapture methods (Liow & Nichols 2010).

## Acknowledgements

The study was funded by the Deutsche Forschungsgemeinschaft (Ko 5382/1-1, Ko 5382/1-2 and Ki 806/16-1) and is part of the Research Unit TERSANE (FO 2332). The authors are grateful to Na Lin for assigning age values to the Cambrian collections. Discussions with M. Steinbauer, E. Jarochowska, J. Pálfy helped in the development of the package, as did the feedback from the first year Master in Paleobiology students of the FAU.

## Author Contributions

ATK conceived the project, wrote the first manuscript draft and the software using code from WK and JA as foundations. WK and CJR contributed to testing, interface and feature development, as well as the debugging of code. All authors contributed to writing the manuscript.

## Data Accessibility

The package is accessible from its GitHub repository (http://github.com/adamkocsis/divDyn) and has been submitted to the CRAN servers. The occurrence data used here are freely available from the Paleobiology Database. All files needed to reproduce the specific example are available on GitHub (http://github.com/adamkocsis/ddPhanero).

